# Multiple brain networks mediating stimulus-pain relationships in humans

**DOI:** 10.1101/298927

**Authors:** Stephan Geuter, Elizabeth A. Reynolds Losin, Mathieu Roy, Lauren Y. Atlas, Liane Schmidt, Anjali Krishnan, Leonie Koban, Tor D. Wager, Martin A. Lindquist

**Author notes:** Authors contributed equally to this work (Tor D. Wager, Martin A. Lindquist). Corresponding author: Stephan Geuter, Department of Biostatistics, Johns Hopkins University, 615 N Wolfe Street, Baltimore, MD 21205, USA.

## Abstract

The brain transforms nociceptive input into a complex pain experience comprised of sensory, affective, motivational, and cognitive components. However, it is still unclear how pain arises from nociceptive input, and which brain networks coordinate to generate pain experiences. We introduce a new high-dimensional mediation analysis technique to estimate distributed, network-level patterns mediating the relationship between stimulus intensity and pain. In a large-scale analysis of functional magnetic resonance imaging data (N=284), we identify both traditional mediators in somatosensory brain regions and additional mediators located in prefrontal, midbrain, striatal, and default-mode regions unrelated to nociception in standard analyses. The whole brain mediators are specific for pain vs. aversive sounds and are organized in five functional networks. Brain mediators explain 32% more within-subject variance of single-trial pain ratings than previous brain-based models. Our results provide a new, broader view of the networks underlying pain experience, as well as distinct targets for interventions.

## Introduction

The brain is central to the generation of pain; it transforms sensory input from peripheral receptors into a complex set of responses, including subjective experience, autonomic and neuroendocrine responses, avoidance behavior, and new learned stimulus-outcome and action-outcome associations. Neurophysiology and neuroimaging studies have identified brain regions that are targeted by afferent nociceptive pathways (Willis and Westlund, 1997; Apkarian et al., 2005; Dum et al., 2009), which are thought to encode sensory-discriminative and affective aspects of pain experience. But pain is a complex experience that entails not only sensory and emotional aspects, but also motivational, attentional and cognitive components. A full picture of the functional brain networks supporting these components of pain experience is still lacking. Here, we address this question using a new multivariate analysis method and a large functional magnetic resonance imaging (fMRI) dataset (N=284).

Although the complexity of pain is widely acknowledged, the underlying brain processes are often conceptualized as a unitary system that is activated by nociceptive input. Brain regions traditionally associated with pain include primary (S1) and secondary (S2) somatosensory, anterior midcingulate cortices (aMCC), medial and lateral thalamus, and posterior and mid-insular cortices (Apkarian et al., 2005; Schweinhardt and Bushnell, 2010; Jensen et al., 2016). But regions not directly targeted by afferent pathways are also activated by acute pain stimuli (Apkarian et al., 2005; Wager et al., 2013; Jensen et al., 2016; Seminowicz and Moayedi, 2017). For example the dorsolateral prefrontal cortex (dlPFC) – a brain region involved in high-level cognitive functions – responds to painful stimulation, shows alterations in chronic pain conditions, and contributes to placebo analgesia (Krummenacher et al., 2010; Bushnell et al., 2013; Seminowicz and Moayedi, 2017; Schafer et al., 2018). Other brain regions, including the ventromedial prefrontal cortex (vmPFC) (Wager et al., 2011; Roy et al., 2012; Geuter et al., 2017b) and the nucleus accumbens (NAc) (Baliki et al., 2012; Chang et al., 2014; Lee et al., 2015; Woo et al., 2015; Ren et al., 2016), key structures for reinforcement learning, also contribute to pain modulation. However, the exact role of these regions is not clear, and they are often thought of as external modulators of activity in the core pain system (Seminowicz and Moayedi, 2017). If so, these regions may be involved in the endogenous construction and regulation of pain in the brain, but they do not mediate the effects of nociceptive input on pain, i.e., they do not link nociception with pain (Woo et al., 2017).

Another view, in line with the ideas originally proposed by Melzack (1999), treats these regions as part of the brain’s pain system for processing cognitive-evaluative aspects of pain. If the neuronal pain system was mirroring the phenomenal complexity of the pain experience, we might expect regions processing cognitive aspects, and perhaps even other areas, e.g., those controlling attention, to be part of the broader pain system (Melzack, 1999). In this case, the dlPFC, vmPFC, NAc, and potentially parietal regions, should be true mediators of the pain response, i.e., they should be closely associated with nociceptive input and pain experience. For example, pain-evoked activity in parietal regions could link attention to the sources of pain with motor intentions (Downar et al., 2003; Oshiro et al., 2007). Furthermore, the different pain mediators should be separable into multiple different functional networks, each associated with different aspects of pain processing.

Here, we introduce a new multivariate mediation analytic framework that captures two important advantages in a single model. First, by analyzing spatial patterns of brain activity, our method builds on spatially distributed information across multiple spatial scales. Second, our method allows the identification of brain responses jointly linked, and interposed between, nociceptive input and pain reports. Mediation analysis (Figure 1A) has previously been applied on a voxel-by-voxel basis to investigate relationships between stimulation intensity, voxel-wise brain activation, and pain report (Figure 1B) (Atlas et al., 2014). However, as with other work on multivariate pattern classification and regression (Wager et al., 2013; Haynes, 2015), a univariate approach can miss brain regions whose contributions to pain perception are conditional on other regions. In order to capture cross-regional interactions, we use a unified high-dimensional approach that takes into account spatial co-variation of activity patterns across the brain (Figure 1C,D).

**Figure 1.**
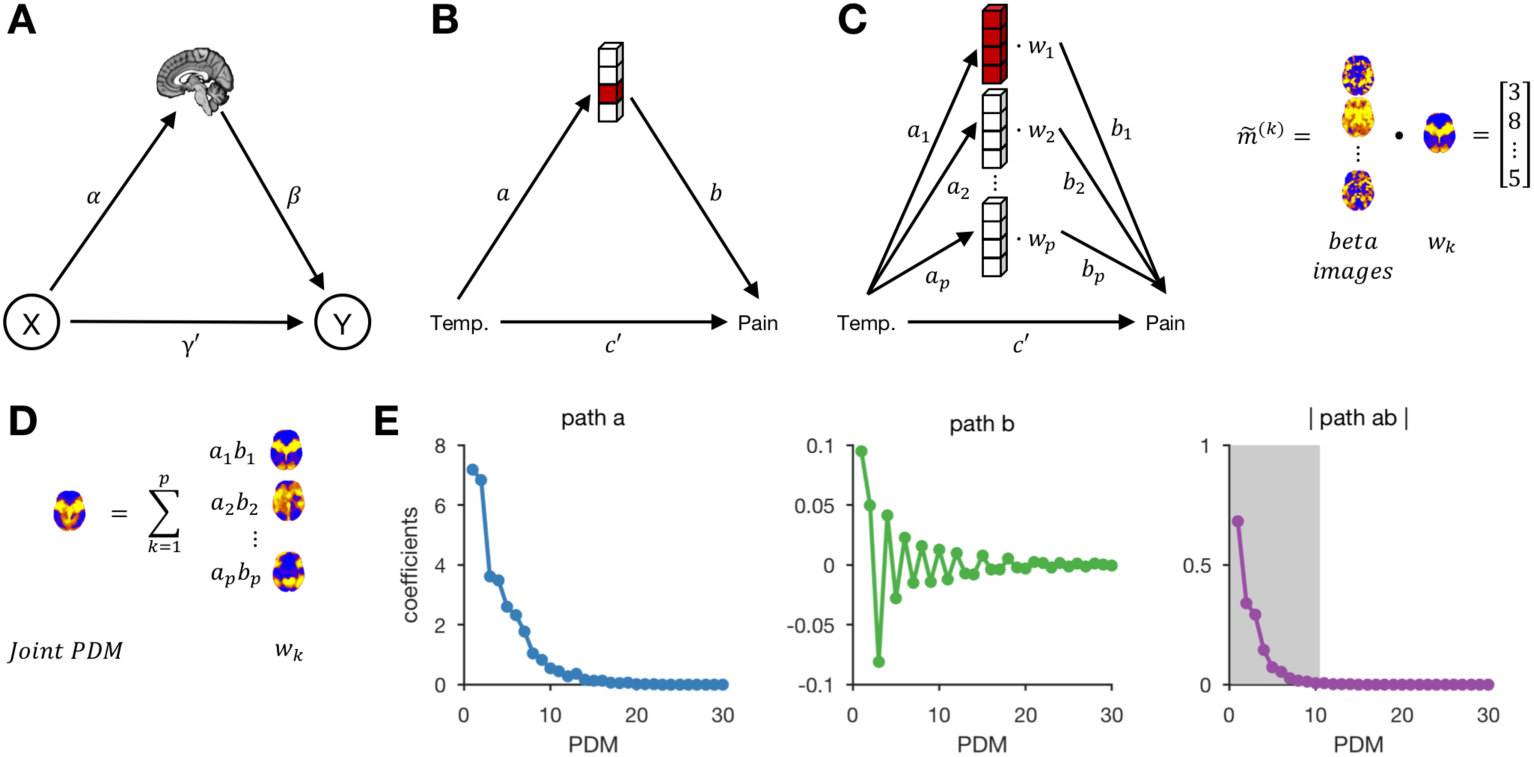
Mediation analysis. (A) Schematic of the mediation analysis framework. Brain activity is an intermediate variable between a manipulated variable *X* and outcome *Y*. (B) In the univariate case, a separate mediation analysis is computed for every brain voxel to determine mediators between stimulation temperature and pain report. (C) In the high-dimensional *Principal Directions of Mediation* (PDM) approach, a linear combination of all brain voxels is used as a mediator. Multiple, orthogonal mediators can be estimated. The weight vectors ***w*_*k*_** (or PDMs) represent the contribution of individual voxels to the ***k*^*th*^** mediation pathway. Taking the dot product of the PDM (***w*_*k*_**) and single-trial brain activation maps (beta images) results in a vector representing a potential mediator. Voxel weights (***w*_*k*_**) are fit so that the indirect, mediated effect is maximal. (D) Individual PDMs can be combined into a single joint PDM by summing the individual PDMs weighted by their path coefficients because individual PDMs are orthogonal to each other. (E) Mediation path coefficients for all 30 PDMs are shown with signs of path ***a*** coefficients set to be positive. Path ***a*** indicates the temperature to brain (PDM) relationship, path ***b*** the PDM to pain rating relationship, and path ***ab*** the indirect, mediated effect. Positive coefficients indicate that voxels with positive weights in a given PDM are positively related with temperature and/or rating. The first 10 PDMs explain more 99% of the total indirect effect. We focus on these PDMs in the following analyses (shaded area in right panel).

This new approach, high-dimensional mediation analysis, identifies multiple whole brain mediators, termed *principal directions of mediation* (PDM). Each PDM represents a pattern of whole brain activity chosen because it maximizes the indirect (mediating) effect between stimulus intensity and pain report. The voxel weights of each PDM inform us about the contribution of individual brain regions to the generation of a painful experience following noxious stimulation. This approach decomposes activity across the brain into multiple networks that independently mediate stimulation effects on outcomes (i.e., pain report). Furthermore, these independent PDMs can be combined into a single, joint PDM that can be prospectively applied to new datasets as a predictive model.

Using data from eight different heat pain studies (N=284), we comprehensively investigate the role of brain mediators in the generation of pain experiences. Seven of the eight studies, were used as training data for the mediation analyses (N=209), and the largest individual study (N=75) was used as a test data set, using the model parameters estimated in the training data to validate model predictions in new individuals. Importantly, the test data set not only included heat pain stimuli, but also physically and emotionally aversive sounds. While brain mediators of pain should generalize to different pain data sets, they are not expected to mediate the relationship between sound stimulation levels and perceived sound intensity. This allows us to study the sensitivity and specificity of the brain mediators of pain.

## Results

### Principal Directions of Mediation (PDM)

Participants in all eight studies underwent fMRI scanning while being exposed to varying levels of heat pain and rating the perceived pain intensity (see Tables 1-3 for details on each study). For each participant, we recorded the temperature applied, the pain rating on a 0-100 scale for each trial and estimated single-trial maps of brain activity. These three variables were used in the primary mediation analysis with temperature as the initial variable, brain activity as the mediator, and pain rating as the outcome variable (Figure 1). Using our novel high-dimensional mediation analysis model (see Chén et al., 2017 for a similar approach), we first estimated 30 whole-brain mediation patterns (PDMs). Each PDM specifies a linear combination of voxels across the brain maximizing the mediated effect from temperature to pain rating, while being orthogonal to other PDMs (Figure 1C). Each PDM (or *w*_*p*_) thus represents a whole brain mediator for pain. For each individual PDM, we obtain path coefficients for the relationship between temperature *X*, brain mediator 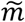, and pain rating *Y* as in a standard mediation model. A positive path *a* indicates that higher temperatures lead to more activity in voxels with positive PDM weights and less activity in voxels with negative PDM weights. A positive path *b* indicates that voxels with positive weights contribute positively to the pain rating after controlling for temperature. This pattern of weights would be expected for regions that receive spinothalamic input, for example the posterior insula or S2 (Willis and Westlund, 1997; Dum et al., 2009), and possibly other mediating regions as well. Finally, we combine the individual brain mediator maps into a joint PDM by computing the weighted sum of the individual PDMs (Figure 1D).

**Table 1.** Demographics

| Study<sup>♦</sup> | Sample Size | Sex | Mean age in Years<br>(Std. Deviation) | Prior publications |
| --- | --- | --- | --- | --- |
| **PDM Training Data** |  |  |  |  |
| **Study 1 (NSF)** | 26 | 9 F / 17 M | 27.8 | Atlas et al. (2014), *PAIN*;<br>Wager et al. (2013) *NEJM* |
| **Study 2 (BMRK3)** | 33 | 22 F / 11 M | 27.9 (9.0) | Woo et al. (2015), *PLoS Biology*<br>Wager et al. (2013) *NEJM* |
| **Study 3 (BMRK4)** | 28 | 10 F / 18 M | 25.2 (7.4) | Krishnan et al. (2016) *eLife* |
| **Study 4 (IE)** | 50 | 27 F / 23 M | 25.1 (6.9) | Roy et al. (2014), *Nature Neuroscience* |
| **Study 5 (ILCP)** | 29 | 16 F* / 12 M | 20.4 (3.3)** | Schmidt et al. (*in prep.*) |
| **Study 6 (EXP)** | 17 | 9 F / 8 M | 25.5 | Atlas et al. (2010), *Journal of Neuroscience* |
| **Study 7 (SCEBL)** | 26 | 11 F / 15 M | 28 (9.3) | Koban et al. (*in prep.*) |
| **PDM Test Data** |  |  |  |  |
| **Study 8 (BMRK5)** | 75 | 39 F / 36 M | 28.2 (5.6) | Losin et al. (*under review*) |

*Note.* <sup>♦</sup>Internal study codes to facilitate tracking of datasets; \*Gender of one participant is unknown; \*\*Age of one participant is unknown. Studies 1-7 have been reported on in Lindquist et al., 2017.

The absolute coefficient values for the indirect *ab* path assess how much of the effect of the manipulated temperature on pain ratings is explained by the brain mediator, i.e., individual PDM pattern. Here, the first 10 PDMs accounted for 99.1% of the total mediation effect (Figure 1E, Figure 1-supplement 1). We thus focus on the first 10 PDMs in all subsequent analyses with minimal loss of information. In order to analyze the contribution of individual brain regions to the mediation of pain, the signs of both paths *a* and *b* and the sign of the voxel weights have to be considered: Voxel weights are multiplied by the respective path coefficients to determine a region’s relationship to stimulation intensity and pain rating. When considering the sign of the voxels weights, four different kinds of relationship are possible: (i) positive to temperature, positive to pain; (ii) negative to temperature, negative to pain; (iii) positive to temperature, negative to pain; and (iv) negative to temperature, positive to pain. Here, type (i) is the standard, positive mediator case and type (ii) represents a negative mediator, in which greater deactivation to the stimulus mediates increased pain (MacKinnon et al., 2000; Atlas et al., 2010). Types (iii) and (iv) are suppressor effects (MacKinnon et al., 2000), e.g., for type (iii), brain activity increases with stimulus intensity that suppress pain, and may thus be involved in stimulus-engaged regulatory processes and other negative feedback loops.

PDM 1 has both positive path *a* and *b* coefficients. Brain regions with positive weights (representing positive mediators, type (i) with positive paths *a* and *b*) are shown in warm colors in Figure 2. These include brain regions commonly associated with pain processing, such as the dorsal posterior and mid-insula, S1, S2, MCC, and the PAG (Figure 2). Significant voxels in MCC stretch into the supplementary motor area (SMA), dorsal of the cingulate sulcus. In addition, PDM 1 contains negative, type (ii), mediators, including the medial prefrontal cortex (mPFC) and ipsilateral S1/M1. The negative weights indicate that these regions show less activation with increasing temperatures and less regional activation is related to higher pain ratings. Such relationships would be expected for brain regions whose function is inhibited by nociceptive input or that are deactivated with increased pain-related processing.

**Figure 2.**
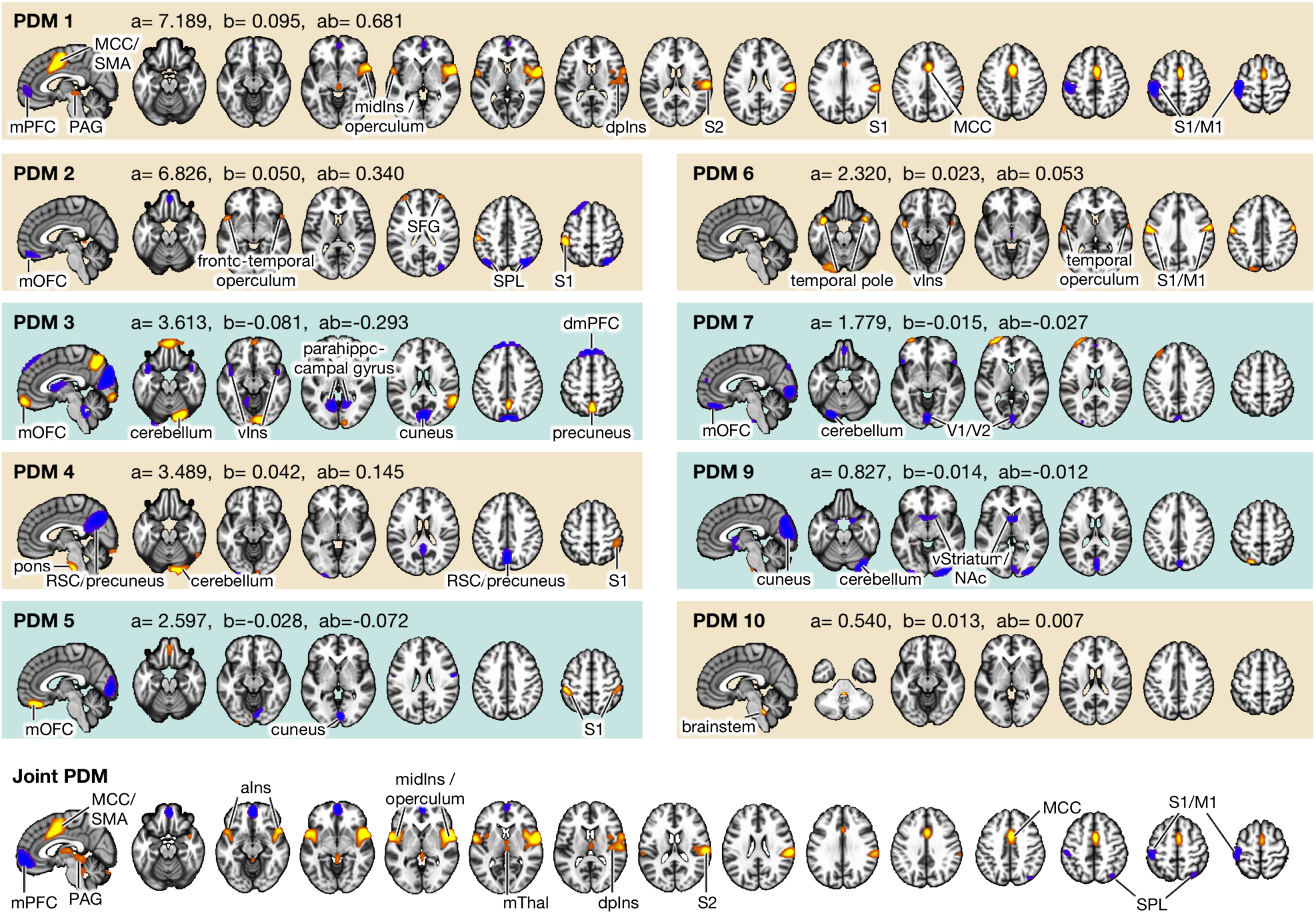
Principal Directions of Mediation. Voxel maps for PDMs with individually significant voxels at *FDR q<0.05*. Tan backgrounds indicate PDMs with positive paths ***a*** and ***b***. Blue backgrounds indicated PDMs with positive path ***a*** and negative path ***b***. Brain activity increases in voxels with positive weights (warm colors) with higher temperatures. Higher brain activity in these voxels is related to higher pain ratings in PDMs with positive path ***b*** (tan panels) and negatively with negative path ***b*** (blue panels). No voxels are individually significant in PDM 8. Bottom panel shows the joint PDM, a combination of the above 10 PDMs. Regions with individually significant voxels in the joint PDM include somatosensory regions, such as S1, S2, insula, MCC, SMA, PAG, and thalamus, but also mPFC M1, and SPL. All brain figures are displayed in neurological convention (left is left) and thresholded at FDR ***q* < 0.05.** MCC=midcingulate cortex, SMA=supplementary motor area, mPFC=medial prefrontal cortex, PAG=periaqueductal gray, midIns=mid-insula, dpIns=dorsal posterior insula, S2=secondary somatosensory cortex, S1=primary somatosensory cortex, M1=primary motor cortex, mOFC=medial orbitofrontal cortex, RSC=retrosplenial cortex, SFG=superior frontal gyrus, vIns=ventral insula, dmPFC=dorsomedial prefrontal cortex, V1=primary visual cortex, V2=secondary visual cortex, vStriatum=ventral striatum, NAc=nucleus accumbens, mThal=medial thalamus, aIns= anterior insula, SPL=superior parietal lobule.

Brain regions positively mediating the relationship between temperature and pain rating (type (i)) in other PDMs are S1, M1, superior frontal gyrus (SFG), fronto-temporal operculum, temporal poles, temporal operculum, ventral insula, pons, and cerebellum (Figure 2). These positive mediators include regions, like the temporal regions, that are traditionally not considered to be pain-processing regions. Brain regions acting as negative mediators (type (ii)) in other PDMs include medial orbitofrontal cortex (mOFC), dorso-medial prefrontal cortex (dmPFC), superior parietal lobule (SPL), retrosplenial cortex (RSC), precuneus, and cuneus.

A more complex function is indicated by positive path *a* coefficients, but negative path *b* coefficients (types (iii) and (iv), PDMs 3,5,7, and 9). Here, regions with positive voxel weights show a positive relationship with temperature, i.e., higher temperatures lead to more activity. By contrast, the negative path *b* indicates that these regions are negatively related to pain ratings controlling for temperature, i.e., more activity is related to lower pain ratings. Regions with such a profile fit a pain-inhibitory role as their activity increases with rising stimulus temperatures, but their increased activity mediates lower pain ratings (type (iii)). Parts of the mOFC, the cerebellum, precuneus, S1, and the left dlPFC fit this pain inhibitory profile.

A final set of regions shows a negative relationship with temperature (positive path *a*, but negative weights) and a positive relationship with pain ratings, controlling for temperature (negative voxel weights and negative path *b* resulting in a net positive relationship; type (iv)). Such regions show stimulus intensity-dependent deactivation, with larger de-activation mediating decreased pain, consistent with regulatory negative feedback mechanisms. Regions with this profile include parts of the mOFC, the parahippocampal gyrus, visual cortices, and the NAc. For example, NAc shows decreased activation for high temperatures, which may relate to punishment or negative reinforcement signals. At the same time, controlling for temperature, stronger NAc de-activation is related to lower pain ratings, potentially signaling reduced motivational relevance.

### Joint PDM

The individual PDMs can be combined into a single, joint PDM since the individual PDMs are orthogonal to each other. Weighting each individual PDM by its indirect effect (path *ab*) and summing the weighted PDMs results in a joint PDM map representing the total contribution of each voxel to the total indirect (pain mediation) effect (see Figure 1D and Methods). Significant voxel weights of the joint PDM map were determined by an additional bootstrap procedure at a false discovery rate (FDR) of *q* < 0.05.

Within the joint PDM, individually significant clusters of positive mediators included S2, MCC, SMA, PAG, insula, including anterior and dorsal-posterior parts, as well as the medial thalamus (Figure 2). Negative mediators (stimulus-induced deactivations mediating increased pain) included mPFC, SPL, S1, and M1. Many of these regions were also part of PDM 1, which accounted for the biggest proportion of the total mediated effect. However, the medial thalamus and SPL were significant in the joint PDM, but not in PDM 1.

While the size of the S1/M1 cluster was smaller in the joint PDM compared to PDM 1, the size of the mPFC cluster increased. Voxel weights for the mPFC and S1 were all negative in the joint PDM. The negative weights in the joint PDM indicate that the net contribution of these regions is a negative mediation of the relationship between temperature and pain, although these regions received positive weights in some of the individual PDMs. Indeed, it is possible for weights to be both positive and negative in different PDMs, because voxels may include neural ensembles participating in different distributed circuits related to either more or less pain. Thus, the individual PDMs represent a decomposition of voxels’ activity into different distributed components, while the joint PDM reflects each voxel’s net contribution (controlling for other voxels). Computing and analyzing the joint PDM can thus help to clarify overall relationships between regional activity and the predictor and outcome variables.

### Clustering PDMs into functional networks

The PDMs provide a dimensional view of coherent distributed processes, with each PDM a distinct dimension; in addition, it is useful to cluster the regions with the highest dimensional weights, to further examine the network structure of the inter-regional relationships. To do this, we used an iterative clustering procedure to group regions based on inter-regional correlations in stimulus-evoked responses across trials without considering stimulation temperatures or pain ratings (Kober et al., 2008; Atlas et al., 2014). The cluster analysis of single-trial activity from significant voxels of all 10 PDMs revealed 26 functional regions organized into 5 different functional networks (Figure 3A,C). A functional description of these networks was determined by computing the similarity of each network with feature maps generated by the meta-analytic tools on neurosynth.org (Yarkoni et al., 2011). The top ten features for each network are shown in Table 4. Network names were chosen based on the functional associations with Neurosynth (Yarkoni et al., 2011) terms. For example, the top three feature associations for network 1 were somatosensory, motor, and stimulation. Based on these associations we labeled network 1 as ‘sensorimotor network’.

**Figure 3:**
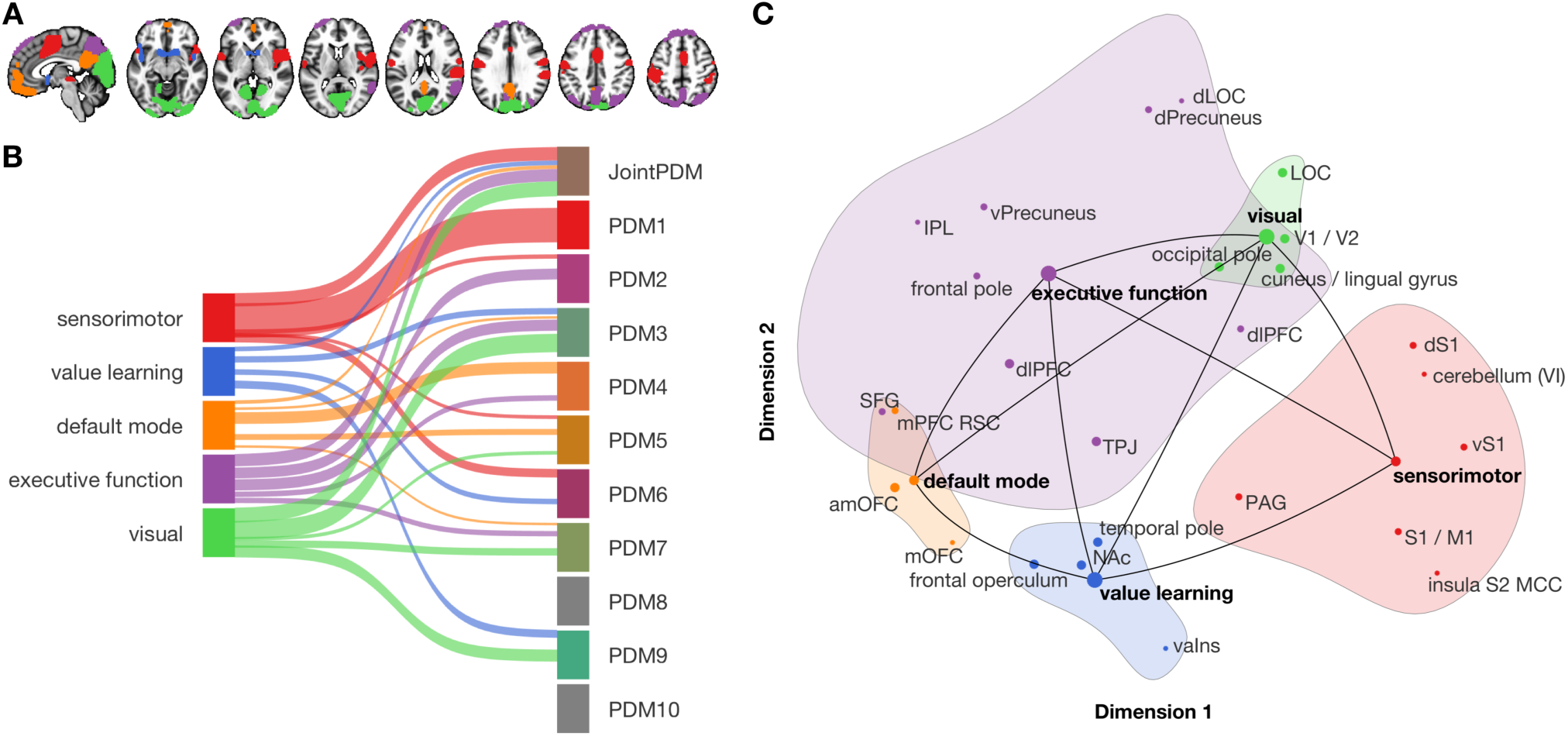
Functional networks mediating pain processing. (A) Five functional networks based on the clustering of brain activity in significant voxels from the PDM analysis. Labels for colors are shown in B. (B) Associations between functional networks and the joint and individual PDMs. Ribbon width represents Dice-coefficient similarity between networks and PDMs. (C) Projection of the five functional networks and individual regions onto the first two dimensions spanned by the NMDS solution. Circle size indicates the number of significant connections for each region or network.

**Table 4.** Neurosynth.org network associations

| **Sensorimotor** |  | **Value-learning** |  | **Default mode** |  | **Executive function** |  | **Visual** |  |
| --- | --- | --- | --- | --- | --- | --- | --- | --- | --- |
| *r* | *features* | *r* | *features* | *r* | *features* | *r* | *features* | *r* | *features* |
| 0.369 | somatosensory | 0.313 | reward | 0.207 | self-referential | 0.139 | mental | 0.223 | visual |
| 0.304 | motor | 0.255 | money | 0.202 | person | 0.124 | intention | 0.152 | eye |
| 0.301 | stimulation | 0.252 | anticipation | 0.201 | self | 0.117 | stories | 0.141 | eyes |
| 0.272 | sensorimotor | 0.252 | rewards | 0.197 | default | 0.115 | attention | 0.137 | color |
| 0.266 | muscle | 0.251 | incentive | 0.176 | autobiographical | 0.115 | visuospatial | 0.126 | shape |
| 0.257 | sensory | 0.240 | monetary | 0.157 | resting state | 0.114 | story | 0.108 | shapes |
| 0.256 | pain | 0.236 | outcome | 0.149 | social | 0.108 | reasoning | 0.105 | spatial |
| 0.245 | movements | 0.196 | outcomes | 0.149 | mentalizing | 0.107 | default | 0.102 | development |
| 0.245 | production | 0.185 | dopamine | 0.148 | personal | 0.106 | calculation | 0.097 | distractor |
| 0.240 | painful | 0.179 | reinforcement | 0.135 | thought | 0.106 | retrieval | 0.097 | target |

*Note:* Top ten features from neurosynth.org showing the highest Pearson's correlation (*r*) with each network.

Network 1 (‘sensorimotor’) included somatosensory regions like dpIns, mid-insula, S2, S1, but also the PAG, MCC, SMA, M1, and cerebellum. The second network (‘value learning’) included the NAc, ventral anterior insula, frontal operculum, and temporal poles. Network 3 consisted of regions that are part of the default mode network (DMN), including mPFC, mOFC, and retrosplenial cortex. The fourth network (‘executive function’) included precuneus, inferior parietal lobule (IPL), superior parietal lobule (SPL), dorsal lateral occipital cortex (dLOC), temporal-parietal junction (TPJ), superior frontal gyrus (SFG), and dlPFC. Finally, network 5 (‘visual’) included mostly occipital, visual areas and parts of the parahippocampal gyrus. The variety of functions ascribed to the five networks mediating pain indicate that pain processing involves multiple, distinct brain networks in addition to somatosensory systems.

We next investigated with which functional networks the individual PDMs are associated by computing pairwise Dice similarity coefficients (Figure 3B). Interestingly, the joint PDM was almost equally similar to the visual (*D* = 0.3), sensorimotor (*D* = 0.27), and to the executive function (*D* = 0.25) networks, again stressing the diversity of brain regions contributing to pain. By contrast, PDM 1 had the greatest overall similarity with any single network, namely with the sensorimotor network (*D* = 0.7). No other network was substantially associated to PDM 1 (all *D* < 0.05). The sensorimotor network was also associated with PDMs 2, 5, and 6. The value learning network was related to PDMs 3, 6, and 9, with the highest similarity to PDM 9 (*D* = 0.16). Similarity between the default mode network and PDM 4 was highest (*D* = 0.24). Parts of the DMN also overlapped with PDMs 3, 5, and 7. The executive function network was associated with PDM 2 (*D* = 0.22) and PDM 3 (*D* = 0.23), and, to a lesser degree, with PDMs 4 and 7. Finally, the visual network was related to PDM 3 (*D* = 0.38) and to a lesser degree to PDMs 5, 7, and 9. The overall similarity pattern between functional networks and PDMs shows that in contrast to PDM1, few of the remaining joint and individual PDMs are dominated by a single network. More often PDMs were comprised of a mix of 2 or 3 networks that together act as a pain mediator, reflecting the complexity of the transformation from nociception into pain experience.

Projecting functional networks and regions onto the first 2 dimensions of the underlying non-metric multidimensional scaling (NMDS) space revealed that the sensorimotor network had high loadings on dimension 1, in contrast to the DMN, which had low loadings on dimension 1 (Figure 3C). Dimension 1 thus spanned from the DMN to the sensorimotor network with the value learning and visual networks located between the two. Dimension 1 could thus be described approximately as an activation-deactivation gradient during pain. Interestingly, the PAG loaded relatively low on dimension 1 within the sensorimotor network and was located closest to the value learning network (Roy et al., 2014). Given the pain-modulatory role of the PAG this could indicate a flexible behavior in pain processing that differs from other regions like S1, S2, or insula, in line with previous literature (Satpute et al., 2013; Roy et al., 2014).

The value learning network loaded lowest on dimension 2. Default mode and sensorimotor networks scored higher than the value learning network, but still lower than the visual network. The executive function network occupied much of the top-left quadrant, with low to medium loadings on dimension 1 and medium to high loadings on dimension 2. With precuneus and LOC as the highest loading regions on dimension 2 and NAc, anterior insula, and mOFC loading lowest, this dimension can potentially be described as a gradient across processing of different time-scales, with NAc encoding transient surprises and precuneus integrating semantic information across longer time-scales (Hasson et al., 2015).

### Validation on an independent cohort

Although we estimated PDMs on a large and diverse data set, here is a risk that the PDMs may over-fit noise inherent in the training data, potentially preventing generalization to other data sets. We thus applied the PDMs to an independent test data set, without re-estimating any model parameters. The resulting vectors of potential mediators 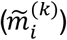 were then entered into standard multi-level mediation models. If the PDMs generalize to the new data, the indirect *a* × *b* effects should be significant on the test data.

Applying the PDMs to independent pain test data (N = 75, an independent community sample cohort of mixed races and sex), revealed significant paths *a* and *b* for all 10 PDMs and the joint PDM (Figure 4A). The indirect path was also significant for the joint PDM and all 10 individual PDMs, suggesting that all 10 PDMs are reliably related to pain and generalize across cohorts. The magnitude of the indirect effects (path *ab*) are monotonically decreasing for the training data (Figure 1E). On the test data, indirect path coefficients were not strictly monotonically decreasing from PDM 1 to PDM 10 (Figure 4A, Figure 4-supplement 1), indicating some variability of the PDM order across data sets, as expected. The joint PDM and the first two individual PDMs had the strongest effect in both data sets, suggesting that they capture the most important brain activity for pain across data sets. Figure 4C shows the predicted pain from the joint PDM plotted against the empirical pain ratings for pain training and test data.

**Figure 4:**
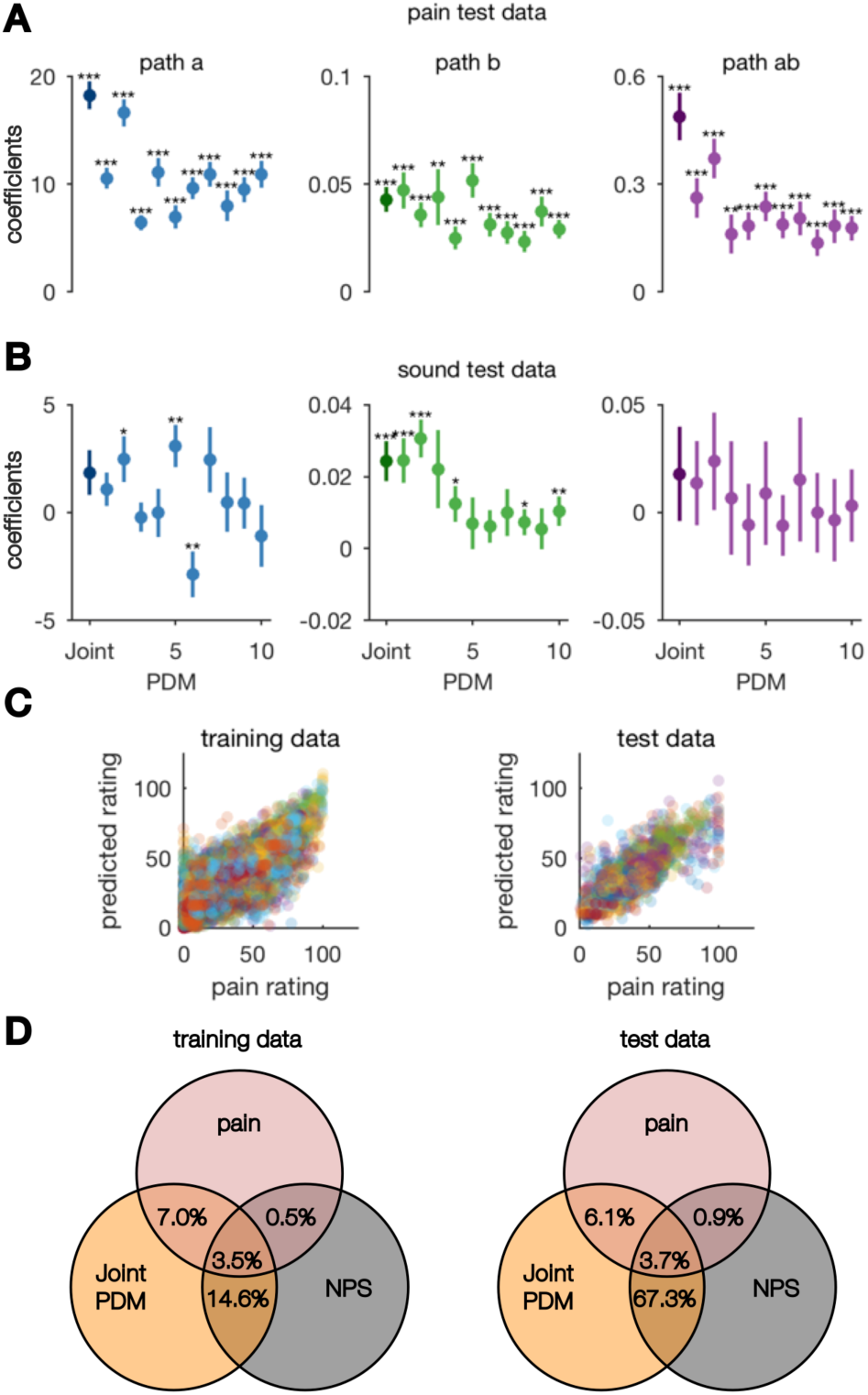
Validation on independent data (N=75). (A) The joint (dark circle) and all 10 individual PDMs (lighter circles) are significant mediators for independent pain test data. (B) PDMs show specificity with respect to aversive sounds because no indirect effect is significant here. (C) Scatter plots of pain predicted from the joint PDM against empirical pain ratings for training (left) and test (right) pain data. Individual trials from all subjects are shown. Colors indicate different subjects. (D) Variance explained in single-trial pain ratings of the training and test data sets for the joint PDM and the NPS, which was only trained on pain ratings without temperature information. The joint PDM accounts for 7% and 6.1%, respectively, pain rating variance not accounted for by the NPS. Error bars indicate SEM. * p<0.05; ** p<0.01; *** p<0.001.

In order to further corroborate the generalizability and robustness of the PDMs, we also estimated 10 PDMs on the original test data set (Study 8) and cross-validated the new PDMs on the original training data set (Studies 1-7). The results were similar to the main results presented here. Six out of ten indirect paths were significant when PDM estimation was done on the smaller sample. The indirect () path coefficients for the first four PDMs were highest when applying the new PDMs to the original training data (Figure 4-supplement 2). Generalization thus does not depend strongly on the choice of the training data.

In order to test whether PDMs are mediators specifically for somatic pain, we also applied the original PDMs to other aversive stimuli that are not painful. The test data of Study 8 also included trials with physically (fingernails on chalkboard) and emotionally (screaming, crying, etc.) aversive sounds with three pre-defined intensity levels of each stimulus type. Study 8 was designed to test specificity vs. generalizability to aversive sounds and matched in duration and approximate aversiveness ratings based on pilot studies; trials were randomly intermixed with heat pain trials. Application of the original PDMs on the sound data revealed no significant indirect effects (Figure 4B, Figure 4-supplement 3) and only nine significant paths *a* or *b* in total. Thus, pain PDMs do not mediate the relationship between sound intensity and intensity ratings for ether type of sound. These results indicate specificity to somatic pain vs. sound.

### Comparison to the Neurological Pain Signature (NPS)

Previous studies have investigated the direct relationship between brain responses and pain reports, both using univariate (Coghill et al., 1999; Bornhövd et al., 2002; Ploner et al., 2010; Atlas et al., 2014) and multivariate approaches (Marquand et al., 2010; Brodersen et al., 2012; Schulz et al., 2012; Wager et al., 2013; Woo et al., 2017). One study trained a multivariate pattern, termed the Neurological Pain Signature (NPS), that predicts pain reports with high accuracy from brain activity that can be easily applied to new data sets (Wager et al., 2013; Krishnan et al., 2016; Geuter et al., 2017a). In contrast to the present approach, the estimation of the NPS did not account for temperature-brain relationships; its goal was to predict pain intensity without demonstrating mediation (Wager et al., 2013). We compared our mediation approach to the predictive power of the NPS by computing the variance explained in single-trial pain ratings by both models. The joint PDM explained a total of 10.5% of the single-trial rating variance in the training data (Studies 1-7) while the NPS explained a total of 4% of the rating variance within subjects (Figure 4D). The variance uniquely explained by the joint PDM was 7%, while the NPS only explained 0.5% unique rating variance and 3.5% of the rating variance was jointly explained by the joint PDM and the NPS. On the test data set (Study 8), the joint PDM explained a unique share of 6.1% of the rating variance. NPS and PDMs explained an additional 3.7% of variance together and the NPS explained additional 0.9% alone (Figure 4D). Together, this indicates that including temperature-brain relationships in the PDM approach captures additional pain variance not explained by the NPS. Here one should note that single-trial data are extremely noisy (Woo et al., 2017), but these numbers indicate high accuracy if an application can average across trials within a person as shown in previous studies results (Wager et al., 2013; Woo et al., 2017).

### Comparison to univariate mediation analysis

In contrast to the present multivariate PDM approach, mass-univariate mediation analyses of fMRI data estimate independent mediation models for each voxel (Wager et al., 2008; Atlas et al., 2014). The intersection of voxels with significant paths *a*, *b*, and *ab* is then interpreted as a set of mediating brain regions. In order to compare the novel high-dimensional PDM approach to the univariate mediation analysis, we first computed a mass-univariate mediation analysis on the training data set (Studies 1-7).

This analysis identified the MCC, cerebellum, posterior and mid-insula, S2, and S1 as brain mediators defined as the intersection of the coefficient maps for paths *a*, *b*, and *ab* at FDR *q* < 0.05 (Figure 5). Comparing these results to the joint PDM, which estimates a joint mediating pattern across voxels, revealed both similarities and some notable differences (Figure 5). The joint PDM included additional regions not in the univariate model, including mPFC, PAG, SPL, and S1. By contrast, the univariate mediation results included a part of the cerebellum that was not included in the joint PDM. Overall, the high-dimensional approach identified more regions than the univariate approach, including regions outside the classic pain processing network like mPFC and SPL. Furthermore, the PAG, a region known to be involved in descending pain control, is part of the joint PDM, but not part of the univariate mediators. Such results are expected if some brain regions make detectable contributions only after controlling for the influences of other brain regions; this is an advantage of multivariate predictive approaches to neuroimaging analysis and multiple regression generally.

**Figure 5:**
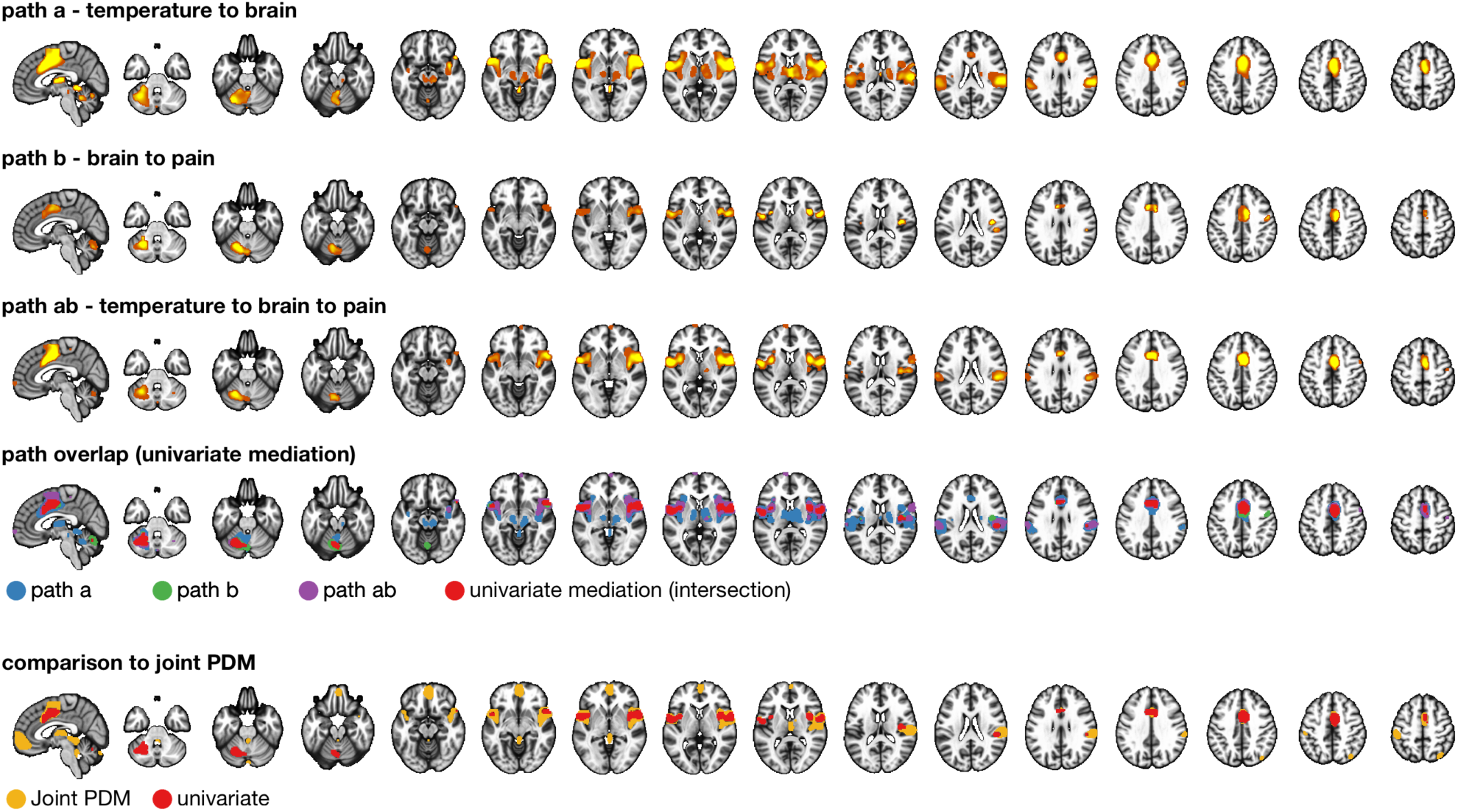
Comparison to univariate mediation analysis. Top three panels show individually significant voxels for paths *a* (blue), *b* (green), and *ab* (purple) from a univariate mediation analysis at FDR *q<0.05*. Panel 4 shows voxels mediating the relationship between temperature and pain, i.e., the overlap between the three paths (red). The bottom panel compares the univariate mediation map (red) and the joint PDM (yellow).

Computing the cosine similarities of PDMs and NPS to both univariate path *a* and *b* maps revealed an interesting pattern (Figure 6). Path *a* represents the relationship between temperature and brain responses, while path *b* represents the relationship between brain responses and rating, controlling for temperature. Projecting all maps on the space defined by temperature and pain rating related brain responses, revealed a linear ordering of components along these dimensions. The map representing the univariate mediation effect (path *ab*) was most similar to path *b* (purple dot in Figure 6). The joint PDM (red) was most similar to the univariate path *a* and located in close proximity to the univariate mediation effect map with respect to the path *a* and *b* maps. However, the cosine similarity of the joint PDM with the univariate path *ab* map was only 0.45, indicating that the two maps reflect substantially different brain processes. Similarities between individual PDMs and the two univariate maps were ordered according to their order of estimation (and the variance explained in the training dataset), with the exception of PDM 3, which was negatively related to both maps. The NPS (black dot) was positioned between PDMs 2 and 4. However, the low overall similarity values for PDMs 3, 5-10, suggest that the space defined by univariate maps of temperature and pain rating related brain responses do not capture all components involved in pain processing. A higher dimensional representation may be more consistent with psychological theories of pain experience.

**Figure 6:**
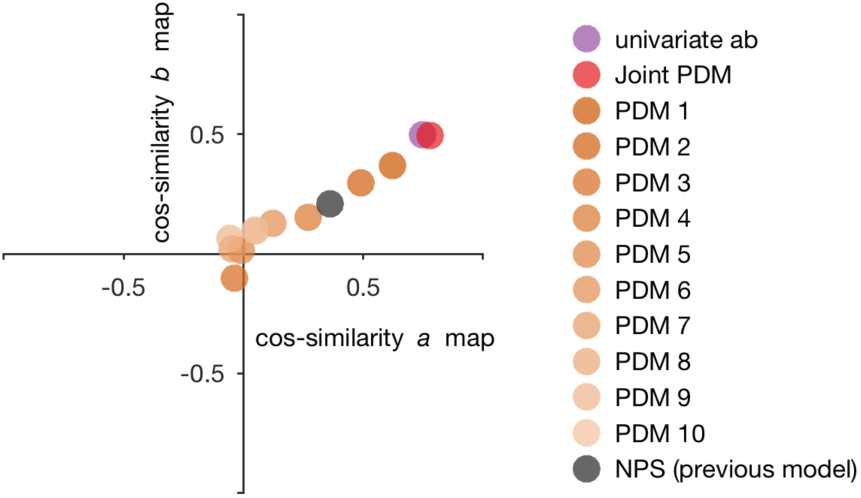
Similarity of mediation maps to univariate stimulus intensity and pain rating maps. Similarity between the PDMs, univariate mediation (path *ab*), and a previous pain predictive map (NPS) to the univariate maps for path *a* and *b*, respectively, measured by the cosine similarity between pairs of maps. The joint PDM and the univariate *ab* map most similar to the *a* and *b* maps representing stimulus intensity and pain intensity, respectively. The similarity between the joint PDM and the path *ab* map is only 0.45. Individual PDMs and the NPS are less and less similar to the univariate effect maps, indicating that the univariate maps do not capture all the information of the multivariate mediation maps.

## Discussion

Using a novel high-dimensional mediation analysis approach (*Principal Directions of Mediation* [PDM]), we identified brain networks that mediate the relationship between stimulus intensity and pain reports. Importantly, the PDM mediators generalized to independent pain test data but not to aversive sound data, suggesting at least some specificity for pain. A parcellation of the brain mediators into functional networks revealed distinct contributions of classic somatosensory brain regions, but also motor regions, value learning, executive control, default mode, and visual regions. This diversity of mediators shows that pain involves many brain regions in addition to somatosensory regions. The observation that the joint PDM map, which integrates the brain mediators, is equally related to the executive function, visual, and sensorimotor networks further supports the importance of non-somatosensory regions in pain processing. In this way, the diversity of brain mediators mirrors the multi-dimensional nature of pain including of sensory, affective, motivational, and cognitive components (Melzack, 1999; Turk and Melzack, 2011).

The new, high-dimensional mediation approach provides a more comprehensive picture of pain processing in the human brain than previous studies using univariate analyses, or studies focusing solely on the stimulation-brain or brain-outcome relationships. This is reflected by the higher share of pain rating variance explained compared to the NPS (Wager et al., 2013), by the higher sensitivity at the brain-voxel level compared to univariate mediation, and by the involvement of brain regions not observed in recent meta-analyses of pain (e.g., mPFC, PAG, and M1) (Duerden and Albanese, 2013; Jensen et al., 2016). The higher sensitivity is demonstrated by the direct comparison of univariate and multivariate maps (Figure 5), and by the PDMs not contained in the space defined by the univariate maps (Figure 6). Together, these results highlight the importance of a broad and methodologically advanced approach to studying pain and related affective processes.

Our results parallel observations in animals and humans that have stressed the importance of psychological and neural processes underlying motivation, learning, attention, and cognition for pain (Melzack, 1999; Atlas et al., 2014; Navratilova and Porreca, 2014; Kucyi and Davis, 2015; Wiech, 2016; Seminowicz and Moayedi, 2017). Functional and structural changes in regions strongly involved in learning, valuation, and executive functions occur during the development of chronic pain and also contribute to chronic pain (Bushnell et al., 2013; Seminowicz and Moayedi, 2017). For example, structural changes observed in the NAc, insula, dlPFC, and sensorimotor cortex distinguish healthy individuals and those suffering from chronic pain (Baliki et al., 2012; Chang et al., 2014; Schwartz et al., 2014; Seminowicz and Moayedi, 2017). Furthermore, altered communication between mPFC and NAc contributes to the development of chronic pain and regulation of acute pain (Baliki et al., 2012; Lee et al., 2015; Woo et al., 2015).

Studies on large-scale functional brain connectivity have also shown that the brain switches dynamically between different states as indexed by state-dependent changes in the communication patterns within and between different brain networks (Cribben et al., 2012; Hutchison et al., 2013). These spontaneous state changes influence perception and cognition (Boly et al., 2007; Sadaghiani et al., 2015) and are known to affect the perception of noxious stimuli (Ohara et al., 2008; Ploner et al., 2010). These observations have led to the hypothesis of a ‘pain connectome’ in which the functional connectivity between networks determines pain experiences (Kucyi and Davis, 2015). The DMN (including mPFC, precunues, and temporal regions), the salience network (including anterior insula, PFC, and TPJ), and the anti-nociceptive network (including mPFC and PAG) have been proposed to be particularly important for pain perception (Kucyi and Davis, 2015). All of these regions were also part of the functional networks mediating pain processing in our multivariate analyses (Figure 3), supporting the notion that pain depends on activation and co-activation patterns (or functional connectivity) between all these regions and not just on the activation level in a unitary core pain system. The high-dimensional mediation approach further allows us to analyze the relationships between activity in individual brain regions with stimulus intensity and pain experience in more detail. In the following we will discuss contributions of brain regions based on their functional relationships with stimulation intensity and pain reports.

Activity in brain regions receiving afferent nociceptive input, including the medial thalamus, PAG, S2, insula, MCC, SMA, and ipsilateral S1 (Dum et al., 2009), increased due to increasing temperatures and higher activity was related to stronger pain, controlling for temperature. This set of commonly pain-associated regions (Apkarian et al., 2005; Bushnell et al., 2013; Duerden and Albanese, 2013; Jensen et al., 2016) was complemented by anterior temporal regions and the cerebellum, which share the same functional response profile. A positive relationship with both temperature and pain rating is in line with a traditional, feedforward encoding view of nociception (Bushnell et al., 2013; Atlas et al., 2014; Geuter et al., 2017a). Because the mediation analysis statistically controls the effects of temperature, our results show that fluctuations in regional activity also contribute to pain perception beyond the regional activity driven by direct afferent input. Activity in these positive mediator regions is thus not only determined by nociceptive input, but the processing and transformation of nociceptive input in these regions contributes to the perceived pain (Büchel et al., 2002).

By contrast, the mPFC, SPL, RSC, precuneus, and contralateral S1 and M1 were negatively related to both temperature and pain. The mPFC, RSC, and precuneus are part of the DMN, which has been associated with mind-wandering and internal thoughts (Andrews-Hanna et al., 2010; Kucyi and Davis, 2015). The negative mediating role of the DMN regions could be related to the disruption of ongoing thought processes by the painful stimulation or attentional refocusing from internal to external sensations. Similarly to the DMN response profile, activity in contralateral M1 was negatively related to stimulus intensity and pain. Motor cortex activity has been associated with painful stimuli in some neuroimaging studies (Apkarian et al., 2005; Schweinhardt and Bushnell, 2010). Along with premotor areas such as SMA, activation in M1 is sometimes interpreted in terms of motor function. However, if M1 activity would represent a motor planning response, we would expect a positive relationship with stimulus intensity and pain ratings. By contrast, the negative relationship of the contralateral M1 with pain is in line with reports of reductions in clinical pain following the inhibition of M1 by transcranial magnetic stimulation (TMS) of M1 (Passard et al., 2007; Mori et al., 2010; Moisset et al., 2016) suggesting a pain modulatory role of M1, potentially via the PAG and the ACC (Pagano et al., 2011). However, further studies are needed to test a potential causal pain inhibitory function of these negative mediator regions in the DMN and sensorimotor cortices.

Pain has also strong motivational implications – humans and animals avoid pain when possible because pain is usually associated with tissue damage (Navratilova and Porreca, 2014; Geuter et al., 2016). It is thus important to learn which stimuli cause pain in order to minimize future harm. However, the role of value learning regions like NAc in pain are complex and not well understood, yet (Becerra et al., 2013; Woo et al., 2015). Unraveling them could contribute a great deal to pain characterization and treatment because of its prominent role in persistent pain in animal models (Chang et al., 2014; Navratilova and Porreca, 2014; Schwartz et al., 2014; Ren et al., 2016) and humans (Baliki et al., 2010, 2012). The present study offers some constraints on interpreting NAc function in pain by demonstrating opposing relationships of NAc activity with stimulus intensity (negative) and pain (positive). Here, NAc shows stimulus intensity-dependent deactivation, with larger de-activation mediating decreased pain, consistent with regulatory negative feedback mechanisms. The NAc might exert its control in this feedback loop indirectly via its connections with the hypothalamus or mPFC as indicated by studies in humans and animals (Baliki et al., 2012; Schwartz et al., 2014; Lee et al., 2015; Woo et al., 2015). However, the exact contribution of the NAc to pain perception might rely on more complex temporal dynamics that cannot be resolved in the current data set. For example, the direction of the valence encoding at pain onset and offset is still a matter of debate (Baliki et al., 2010; Becerra et al., 2013) as is its role in aversive learning more generally (Roy et al., 2014; Matsumoto et al., 2016). Elucidating the specific contributions of the NAc in different contexts in future studies will further help our understanding of motivational and learning aspects for pain perception.

An advantage of the present multivariate mediation approach is that it controls for the effects of stimulation intensity when estimating the relationship between brain activity and reported outcomes. Compared to approaches that do not take into account the stimulus-brain relationship when predicting pain (Wager et al., 2013; Krishnan et al., 2016; Lindquist et al., 2017), the present mediation approach yields higher predictive accuracy. Both approaches may yield whole brain maps that can be used as predictive models of acute pain that can be applied prospectively to new data. Application of the PDMs to new datasets can be used to (i) further evaluate the sensitivity and specificity of the model, and (ii) to evaluate the effects of psychological or medical interventions on the brain processes supporting pain. Testing the PDMs on a large, independent data set showed that the PDMs generalize to other pain data, but do not generalize to aversive sounds. This speaks against the notion that the brain mediators are completely driven by stimulus independent features, such as general feelings of aversiveness or unpleasantness.

In summary, the new high-dimensional mediation analysis revealed a comprehensive picture of brain responses underlying the complex, multi-faceted pain experience. Several brain regions, such as the mPFC, NAc, and M1, are shown to directly and formally mediate stimulus-to-pain relationships. The functional diversity of the brain mediators observed here offers a better understanding of the brain responses underlying the complexity of the pain experience.

## Acknowledgements

We are grateful for the funding support of the DFG (GE 2774/1-1 to S.G.) and the NIH, which supported this work under grants R01DA035484 (T.D.W.), 2R01MH076136 (T.D.W.), R01DA027794 (T.D.W.), R01 EB016061 (M.A.L.) and P41 EB015909 (M.A.L.).

## Author Contributions

S.G., T.D.W, and M.A.L. designed the study. M.A.L. contributed unpublished analytical tools. S.G. conducted data analysis and drafted the manuscript. S.G., T.D.W., M.A.L., and E.A.R.L. edited and revised the manuscript. E.A.R.L., M.R., L.Y.A., L.S., A.K., and L.K. curated neuroimaging data and provided comments on the manuscript.

## Materials and Methods

### Participants

The analysis included a total of 284 healthy participants from 8 independent studies, with sample sizes ranging from N = 17 to N = 75 per study. Descriptive statistics on the age, sex, and other features of the subjects in each individual study are provided in Tables 1-3. Further details on Studies 1-7, which were used to estimate the PDMs are provided in Lindquist et al. (2017). Participants were recruited from New York City and Boulder/Denver Metro Areas. The institutional review board of Columbia University and the University of Colorado Boulder approved all the studies, and all participants provided written informed consent. Preliminary eligibility of participants was determined through an online questionnaire, a pain safety screening form, and a functional Magnetic Resonance Imaging (fMRI) safety screening form.

We applied several exclusion criteria for analysis purposes. Participants with psychiatric, physiological or pain disorders, neurological conditions, and MRI contraindications were excluded prior to enrollment. In addition, participants were required to have at least 30 trials with low variance inflation factors (see below), non-missing rating, and stimulation intensity data. Based on these criteria, 18 participants from Study 8 were excluded, resulting in a total of 209 participants for the primary PDM analysis and 75 participants for the validation sample.

### Procedures

In all studies, participants received a series of contact-heat stimuli and rated their experienced pain following or during each stimulus. The number of trials, stimulation sites, intertrial intervals, rating scales, and stimulus intensities and durations varied across studies, but were comparable; these variables are summarized in Tables 2 and 3. Each study also comprised a specific psychological manipulation (except Study 8), such as placebo treatment, which will be or has been reported elsewhere (Table 1). Study 8, which was used for validation purposes (see below), also presented aversive sounds to participants. Trials with aversive sounds were used to test the specificity of the pain PDMs. Sounds included were a physically aversive recording of nails on a chalkboard and a set of emotionally aversive sounds (attacks, screaming, and crying) from the International Affective Digital Sounds database (IADS) (Bradley and Lang, 2007). Aside from these sound trials, we focus on brain mediation of pain across all trials in the present paper, irrespective of the study-specific psychological and physical manipulations that influenced pain.

**Table 2.** Stimulation Parameters

| Study | Intensities | Mean Temperature by Intensity Level<br>(Within Subject SE) | Rating scale | Mean Ratings by Intensity Level<br>(Within Subject SEM) |
| --- | --- | --- | --- | --- |
| **PDM Training Data** |  |  |  |  |
| **Study 1**<br>**(NSF)** | N, L, M, H<br>(Calibrated) | 40.8, 43.1, 45.1, 47.0<br>(0.16) | 0-8 VAS (0, no sensation; 1, non-painful warmth; 2, low pain; 5, moderate pain; 8, maximum tolerable pain) | 2.0, 2.8, 4.2, 6.6<br>(0.14) |
| **Study 2**<br>**(BMRK3)** | 6 levels<br>(Fixed) | 44.3, 45.3, 46.3, 47.3, 48.3, 49.3 | 0-100 VAS | 49.1, 56.6, 74.3, 99.4, 133.0, 159.3<br>(3.12) |
| **Study 3**<br>**(BMRK4)** | L, M, H<br>(Fixed) | 46.0, 47.0, 48.0 | 0-100 VAS (0, no sensation; 1.4, barely detectable; 6.1, weak; 17.2, moderate; 35.4, strong; 53.3, very strong; 100, strongest imaginable sensation) | UL: 31.7, 40.5, 53.6<br>(0.9787)<br>LL: 31.5, 40.2, 53.3<br>(0.96) |
| **Study 4**<br>**(IE)** | L, M, H<br>(Fixed) | 46.0, 47.0, 48.0 | 0-100 VAS (0, no pain; 100, worst imaginable pain) | 29.4, 38.9, 51.9<br>(0.64) |
| **Study 5**<br>**(ILCP)** | L, H<br>(Calibrated) | 44.7, 46.7<br>(0) | 0-8 VAS (no pain to worst pain imaginable) | 24.3, 46.7<br>(1.14) |
| **Study 6**<br>**(EXP)** | L, M, H<br>(Calibrated) | 41.2, 44.4, 47.2<br>(0.21) | 0-8 VAS (0, no sensation; 1, non-painful warmth; 2, low pain; 5, moderate pain; 8, maximum tolerable pain) | 2.5, 4.3, 7.4<br>(0.13) |
| **Study 7**<br>**(SCEBL)** | L, M, H<br>(Fixed) | 48, 49, 50 | 0-100 VAS (0, no pain; 100, worst imaginable pain) | 26.0, 33.3, 40.4<br>(1.12) |
| **PDM Test Data** |  |  |  |  |
| **Study 8**<br>**(BMRK5)** | L, M, H<br>(Fixed) | 47, 48, 49 | 0-100 gVAS (0, no experience; 100, strongest imaginable experience) | 30.6, 39.9, 48.2<br>(1.64) |

*Note:* Heat /pain levels: N = Nonpainful, L = Low, M = Medium, H = High. VAS = visual analogue scale. gVAS = generalized visual analogue scale.

**Table 3.** Task Characteristics

| **Study** | **Duration**<br>**(seconds)** | **Inter-heat**<br>**interval**<br>**(seconds)** | **Locations**<br>**(number of**<br>**sites)** | **Range of**<br>**Number of Trials**<br>**Per Subject** | **Mean proportion of**<br>**trials excluded**<br>**(Std. Deviation)** | **Other experimental**<br>**manipulations** |
| --- | --- | --- | --- | --- | --- | --- |
| **PDM Training Data** |  |  |  |  |  |  |
| **Study 1**<br>**(NSF)** | 10 | 38 | Left arm (3) | 35-48 | 0.08 (0.07) | Masked emotional faces evenly<br>crossed with temperature |
| **Study 2**<br>**(BMRK3)** | 12.5 | 20.5-28.5 | Left arm (2) | 97 | 0.1 (0.04) | Cognitive self-regulation up and<br>down |
| **Study 3**<br>**(BMRK4)** | 11 | 25-27 | Left arm (4),<br>left foot (4) | 81 | 0.08 (0.06) | Heat-predictive visual cues (low,<br>medium, or high) |
| **Study 4**<br>**(IE)** | 11 | 36-38 | Left arm (6) | 48 | N/A | Heat-predictive visual cues;<br>placebo manipulation |
| **Study 5**<br>**(ILCP)** | 10 | 17-25 | Left arm (2) | 64 | 0.05 (0.03) | Agency (make choice, observe<br>choice), Certainty (80% low pain,<br>50% low pain) |
| **Study 6**<br>**(EXP)** | 10 | 38 | Left arm (4) | 61-64 | 0.03 (0.04) | Heat-predictive auditory cues |
| **Study 7**<br>**(SCEBL)** | 1.85 | 26-37 | Right leg (6) | 96 | 0.04 (0.03) | Heat-predictive visual cues (low or<br>high) and unreinforced social<br>information |
| **PDM Test Data** |  |  |  |  |  |  |
| **Study 8**<br>**(BMRK5)** | 8, 11 | 11.5-32.75 | Left arm (4) | 30-36 | 0.04 (0.04) | Aversive sounds, modality-<br>predictive cues (sound vs. heat) |

### Thermal stimulation

In each study, except Studies 7 and 8, thermal stimulation was delivered to multiple skin sites using a TSA-II Neurosensory Analyzer (Medoc Ltd., Chapel Hill, NC) with a 16 mm Peltier thermode endplate. A PATHWAY system (Medoc Ltd., Chapel Hill, NC) was used in Studies 7 and 8. Study 7 used a circular CHEPS Peltier endplate (diameter: 32 mm) and study 8 used a 16 mm ATS Peltier endplate. On every trial, after the offset of stimulation, participants rated the magnitude of the warmth or pain they had felt during the trial on a visual analog scale. Participants in Study 8 rated their pain continuously during stimulation. The maximum rating of each trial was used in the following analyses. Other thermal stimulation parameters varied across studies, with stimulation temperatures ranging from 40.8 °C to 50 °C and stimulation durations from 1.85 to 12.5 s. Most studies applied thermal stimulation to the forearm. See Table 2 for stimulation intensity levels, mean temperature for each intensity level, and details of the rating scales. See Table 3 for stimulation duration, duration of inter-stimulus interval, number and location of stimulation sites, and number of trials per subject.

### fMRI data processing

#### Preprocessing

Structural T1-weighted images were co-registered to the mean functional image for each subject using the iterative mutual information-based algorithm implemented in SPM (Ashburner and Friston, 2005), and then normalized to MNI space using SPM. The version of SPM used varied across studies (Studies 1 and 6 used SPM5; while all other studies used SPM8; http://www.fil.ion.ucl.ac.uk/spm/). Following normalization, Studies 1 and 6 included an additional step of normalization to the group mean using a genetic algorithm-based normalization (Wager and Nichols, 2003; Atlas et al., 2010, 2014).

For each functional dataset, initial volumes were removed to allow for image intensity stabilization (see Lindquist et al. (2017) for details). In addition, volumes with signal values that were outliers within the time series (i.e., “spikes”) were removed. To identify outliers, both the mean and the standard deviation of intensity values across each slice were computed for each image. The Mahalanobis distances for the matrix of (concatenated) slice-wise mean and standard deviation values by functional volumes (over time) were computed, and values with a significant *X*^2^ value (corrected for multiple comparisons based false discovery rate) were considered outliers. In practice, less than 1% of images were deemed outliers. The output of this procedure was later included as nuisance covariates in the subject level models. Next, functional images were corrected for differences in the acquisition timing of each slice (except for multiband data with a short TR of 480 ms in Study 8) and were motion-corrected (realigned) using SPM. The functional images were warped to SPM’s normative atlas (warping parameters estimated from co-registered, high-resolution structural images), interpolated to 2 × 2 × 2 *mm*^3^ voxels, and smoothed with an 8 mm FWHM Gaussian kernel.

#### Single trial analysis (Except Study 3 and Study 6)

For each study, a single trial, or “single-epoch”, design and analysis approach was used to model the data. Quantification of single-trial response magnitudes was done by constructing a GLM design matrix with separate regressors for each trial (Rissman et al., 2010; Mumford et al., 2012). First, boxcar regressors, convolved with the canonical hemodynamic response function (HRF), were constructed to model cue and rating periods in each study. Regressors for each trial, as well as several types of nuisance covariates were also included. Because each trial consisted of relatively few volumes, trial estimates could be strongly affected by acquisition artifacts that occur during that trial (e.g. sudden motion, scanner pulse artifacts, etc.). Therefore, trial-by-trial variance inflation factors (VIFs; a measure of design-induced uncertainty due, in this case, to collinearity with nuisance regressors) were calculated, and any trials with VIFs exceeding 2.5 were excluded from the analyses (VIF threshold for Study 8 was 3.5 as in the primary publication). For Study 1, global outliers (trials that exceeded three standard deviations (SDs) above the mean) were also excluded, and a principal component based denoising step was employed during preprocessing to minimize artifacts. This generated single trial estimates that reflect the amplitude of the fitted HRF on each trial and refer to the magnitude pain-period activity for each trial in each voxel.

#### Single trial analysis (Only Study 3 and Study 6)

For Studies 3 and 6, single trial analyses were based on fitting a set of three basis functions, rather than the standard canonical HRF used in the other studies. This flexible strategy allowed the shape of the modeled hemodynamic response function (HRF) to vary across trials and voxels. This procedure differed from that used in other studies because it maintains consistency with the procedures used in the original publications. For both Study 3 and Study 6, the pain period basis set consisted of three curves shifted in time and was customized for thermal pain responses based on previous studies (Lindquist et al., 2009; Atlas et al., 2010). To estimate cue-evoked responses for Study 6, the pain anticipation period was modeled using a boxcar epoch convolved with a canonical HRF. This epoch was truncated at 8 s to ensure that fitted anticipatory responses were not affected by noxious stimulus-evoked activity. As in the other studies, nuisance covariates were included and trials with VIFs larger than 2.5 were excluded. In Study 6 trials that were global outliers (those that exceeded 3 SDs above the mean) were also excluded. The fitted basis functions from the flexible single trial approach were used to reconstruct the HRF and compute the area under the curve (AUC) for each trial and in each voxel. These trial-by-trial AUC values were used as estimates of trial-level pain-period activity.

#### Data sets and PDM validation

The high-dimensional brain mediators (PDMs, see below) were estimated on the training data comprised of Studies 1-7 (Lindquist et al., 2017). Even though this data set is large (N=209) and diverse, the possibility of overfitting in the training data might reduce the generalizability of the PDMs. To test for the generalizability of the PDMs, we validated the PDMS on independent test data (Study 8, N=75). Computing the inner product of each PDM with each single-trial beta image from Study 8 resulted in 10 potential mediator variables. Each of these potential mediators was then subjected to a multi-level mediation analysis (Wager et al., 2009) with *p*-values determined by a bootstrap procedure with 5,000 iterations each. If the PDMs generalize to the new dataset, paths a*_k_*, b*_k_*, and the indirect effect ab_*k*_ should be significant for all *k* = 1, …, 10 PDMs.

We also tested whether the PDMs specifically mediate the relationship between temperature and pain intensity. To this end, we also tested the original PDMs on the aversive sound trials from Study 8. If the PDMs reflect specific patterns of brain activity involved in pain processing, they should not mediate the relationship between sound stimulation level and intensity ratings. We thus expect no significant indirect effect for the sound trials.

A further test to validate the stability of PDM estimation was conducted by switching training and test data. That is, pain PDMs were estimated on Study 8 and tested on the original training data from Studies 1-7 as described above.

#### Dimension reduction

The training data set consisted of a total of 13,372 single-trial beta images, each consisting of 229,519 voxels, from 209 participants. To reduce the dimensionality of the data to a computationally tractable size, a generalized version of population value decomposition (PVD) (Caffo et al., 2010; Crainiceanu et al., 2011; Chén et al., 2017) was applied (using PVD.m as part of the M3 mediation toolbox available at https://github.com/canlab/MediationToolbox). This procedure is similar to singular value decomposition (SVD) but decomposes the data matrix into both participant specific and population specific components. We chose a dimensionality of *p* = 30 based on a tradeoff between variance explained and the number of trials available for each participant. The beta images were z-scored within each participant before PVD application. The reduced data matrix used for Principal Directions of Mediation (PDM) estimation consisted of a matrix with dimensions 13, 372 × 30.

### Principal Directions of Mediation (PDM)

Let X*_i_* be the temperature, Y_*i*_ the reported pain, and 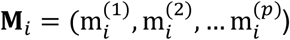 the brain activity over *p* voxels (i.e., the beta maps) measured between the application of the thermal stimuli and the pain report for observation (i.e., trial) *i* = 1, … *n*. We are interested in determining how brain activation mediates the relationship between temperature and pain report, which is illustrated using the three-variable path model shown in Figure 1. We can estimate the parameters of this model using the following set of equations:

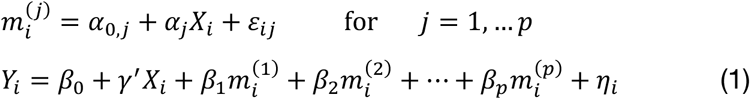

Once the parameters have been estimated we can express the total effect W as the sum of the direct and indirect effects as follows:

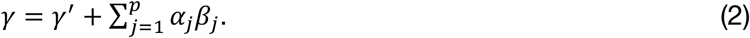

If *p* is relatively small the series of regressions described in (1) can be used to estimate the pertinent mediation effects. However, in our setting there are too many mediators to allow reasonable interpretation (unless the model coefficients are highly structured) and there are many more mediators than subjects, precluding estimation using standard procedures. To overcome these problems, we introduce a transformation of the space of mediators, determined by finding linear combinations of the original mediators that (i) are orthogonal; and (ii) are chosen to maximize the indirect effect. The first constraint allows us to fit a separate linear model for each transformed variable. The second constraint allows us to limit our analysis to only those directions that contain the most information about the indirect effect. Here, we improve and extend the approach proposed by Chén et al. (2017) by choosing a different cost function, computing the joint PDM, and analyzing an almost 10-times larger data set.

This new model, called the *principal directions of mediation* (PDM), linearly combines activity in different voxels into a smaller number of orthogonal components, with components ranked based upon the proportion of the indirect effect that each accounts for. Ideally, the components form a small number of uncorrelated mediators that represent interpretable networks of voxels.

To illustrate, let 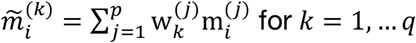 be a set of linear transformations of the mediators with 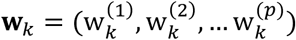. Placing these new variables into our mediation model we obtain:

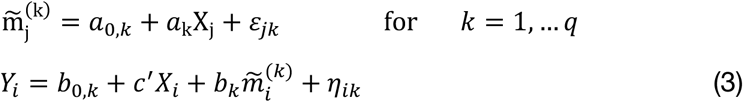

Now, we can decompose the total effect into direct and indirect effects as follows:

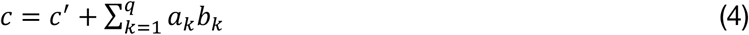

The difference between this model and the standard mediation model described in (1) is that the **w***_k_* are unknown. In our approach **w**_1_ is chosen so that it maximizes the amount of the indirect effect that is explained (i.e., a_1_b_1_ is maximized). We refer to **w**_1_ as the first *principal direction of mediation* (PDM). Note the first PDM corresponds to voxel-specific weights that can be mapped onto the brain, and thus provides interpretable maps of brain networks in the same manner as independent component analysis (ICA) and principal component analysis (PCA). Subsequent directions **w***_k_*, *k* = 1, … *q*, can be found that maximize the remaining indirect effect conditional on being orthogonal to previous PDMs. As the transformed mediators are ranked based upon the proportion of the indirect effect explained, one could potentially limit the number of PDMs computed to achieve dimension reduction. Hence, our approach is philosophically similar to PCA, but addresses a fundamentally different problem.

The individual, orthogonal PDMs can be combined into a joint PDM by computing the following weighted sum:

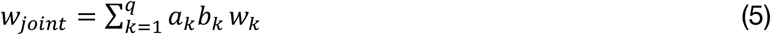

According to the model formulation the signs of the PDMs are not identifiable, as any change in the sign of 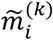 can be offset by a change in sign of both a_*k*_ and b_*k*_. We fix the signs of a_*k*_ to be positive for easier interpretation, i.e., positive voxel weights indicate higher brain activity for higher stimulus intensities. This is a similar constraint to the ICA approach often used in neuroimaging to detect networks. Note this does not impact the joint PDM as the sign of a*_k_*b*_k_*is unchanged if both a_*k*_ and b_*k*_ change signs.

The problem of finding the *k*^*th*^ PDM involves finding the vector **W**_*k*_ that maximizes a*_k_*b*_k_* based on the constraint that 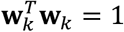 and 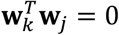 for all *j* = 1, …, *k* − 1. This problem can be solved using a nonlinear programming solver such as the interior-point algorithm. Inference is performed using a bootstrap procedure with 5,000 iterations, as described in Chén et al. (2017). We also test individual voxel weights for the joint PDM for significance using the bootstrap procedure above. All PDM maps are thresholded at a false discovery rate (FDR) of *q* < 0.05. We present results of 10 PDMs accounting for more than 99% of the total indirect effect (Figure 2). The PDM implementation is available at https://github.com/canlab/MediationToolbox (multivariateMediation.m).

In summary, we obtain scalar coefficients for paths a_*k*_, b_*k*_, and c′_*k*_, as well as the indirect effect (): for each PDM as in a standard, univariate mediation analysis. In addition, we obtain the voxel weight vector **w**_*k*_ that maximizes the indirect effect *ab*_*k*_.

### Cluster analysis

The voxel weight maps for the mutually independent 10 PDMs span a high-dimensional space of brain mediators of pain perception. In order to reduce the dimensionality of that space and identify brain regions with similar activation profiles, we conducted a two-stage cluster analysis. The procedure is described in detail in Kober et al. (2008) and Atlas et al. (Atlas et al., 2014). Briefly, for significant voxels from the 10 PDMs we extracted single-trial activity estimates, resulting in a 13,372 trials × 25,469 voxels matrix. We then used singular value decomposition (SVD) to reduce the dimensionality of the voxel space. We kept 364 components that explained 95% of the variance. Next, we clustered voxels into 250 parcels using hierarchical clustering. We then computed average single-trial activity within each parcel and used non-metric multidimensional scaling (NMDS) and hierarchical clustering to further reduce the dimensionality of the data. Inspection of the Shepard plot suggested a NMDS dimensionality of 15 with stress indices below 0.05. Stress indices (j) are computed according to Shepard (1980) with

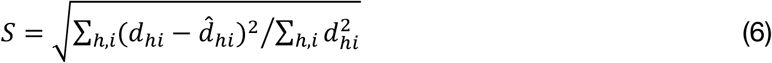

Here, *d*_*hi*_ is the pairwise empirical dissimilarity and 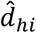 is the distance implied by the current solution between two brain regions ℎ and K. Hierarchical clustering was then used to cluster the 250 parcels into 33 regions that co-activate across trials. These regions were not necessarily contiguous and some spanned multiple anatomical regions, e.g., covering right mid-, and dorsal insula plus operculum. Since we used voxel-wise FDR correction on the 10 PDMs, we expect some false positive values. Accordingly, some of the functional regions were located in the cerebrospinal fluid or outside the gray matter. We thus removed 7 smaller functional clusters that were considered highly unlikely to be true gray matter region. We then averaged brain activity within the remaining 26 functional regions. NMDS was used to reduce the dimensionality again to 10 dimensions based on stress values. Applying hierarchical clustering again on the regions identified in the previous step identified large-scale functional brain networks. Permutation tests indicated that 5 networks provided the best clustering solution in terms of improvement over solutions on permuted data. The position of the 5 networks and their constituent brain regions were projected on the first 2 dimensions of the NMDS space to visualize relationships and functional connectivity. Similarity of those 5 networks with the binarized PDM maps was assessed by Dice coefficients, which represents the true positive rate of the intersection between two maps.

### Univariate mediation analysis

In univariate mediation analyses, a mediation model is estimated separately for every brain voxel (Wager et al., 2008; Atlas et al., 2010, 2014). Univariate mediation analysis produces three sets of brain maps – one for each path – in contrast to the PDM approach, which estimates only one set of paths for each PDM map. Previous studies also used smaller sample sizes available than the present study and had thus less statistical power than the present study. We ran a univariate mediation analyses on the training data set to directly compare the univariate results to the PDM approach. Univariate multilevel mediation analysis was conducted using the Multilevel Mediation and Moderation (M3) Toolbox for Matlab (https://github.com/canlab/MediationToolbox). Voxel-wise significance was determined using a bootstrap procedure with 5,000 iterations. A false discovery rate (FDR) of *q* < 0.05 was used to control for multiple comparisons.

---

**Figure 1-supplement 1.**
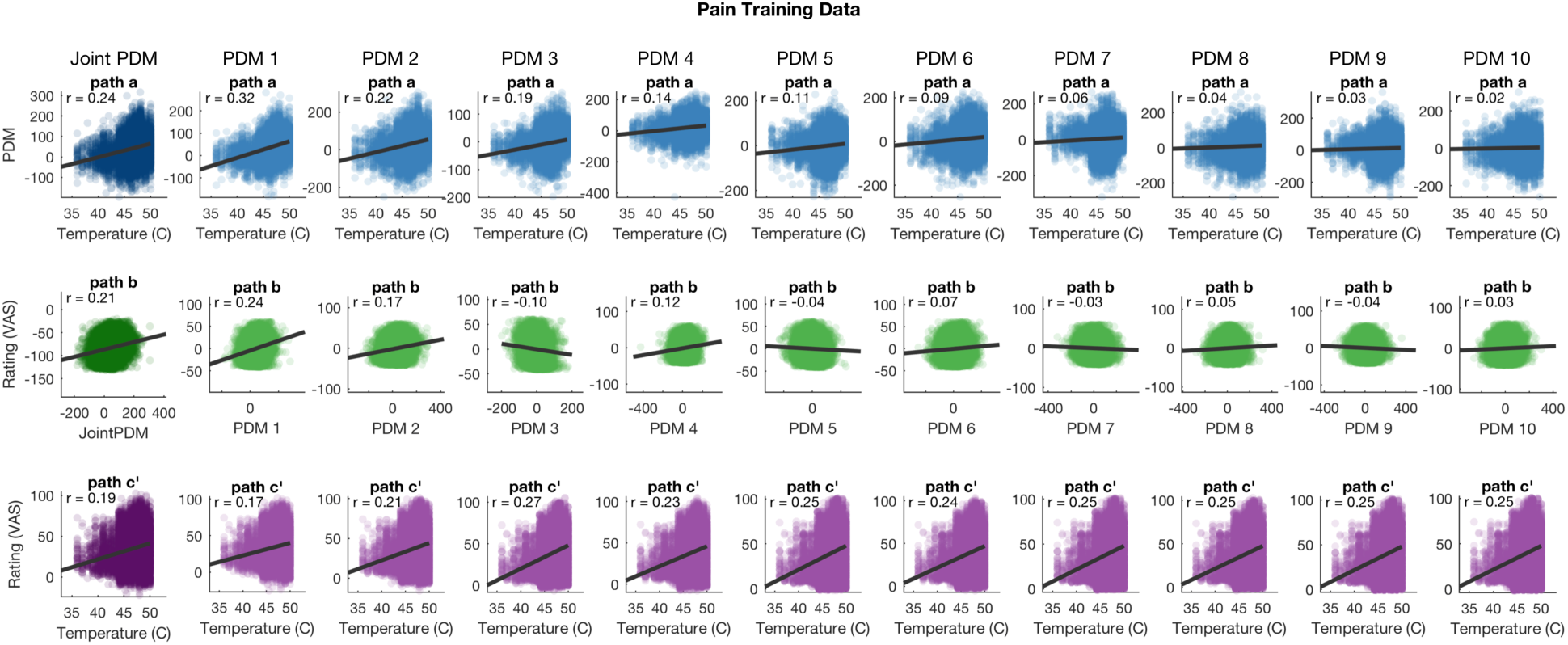
Bivariate relationships between temperatures, mediators (PDM expression), and pain ratings for the training data (studies 1-7). Data are adjusted according to the mediation equations, i.e. ratings in path b plots are adjusted for temperatures and PDMs, ratings in path c’ plots are adjusted for PDMs, and PDMs in path b plots are adjusted for temperatures. PDMs are estimated on the training data.

**Figure 4-supplement 1.**
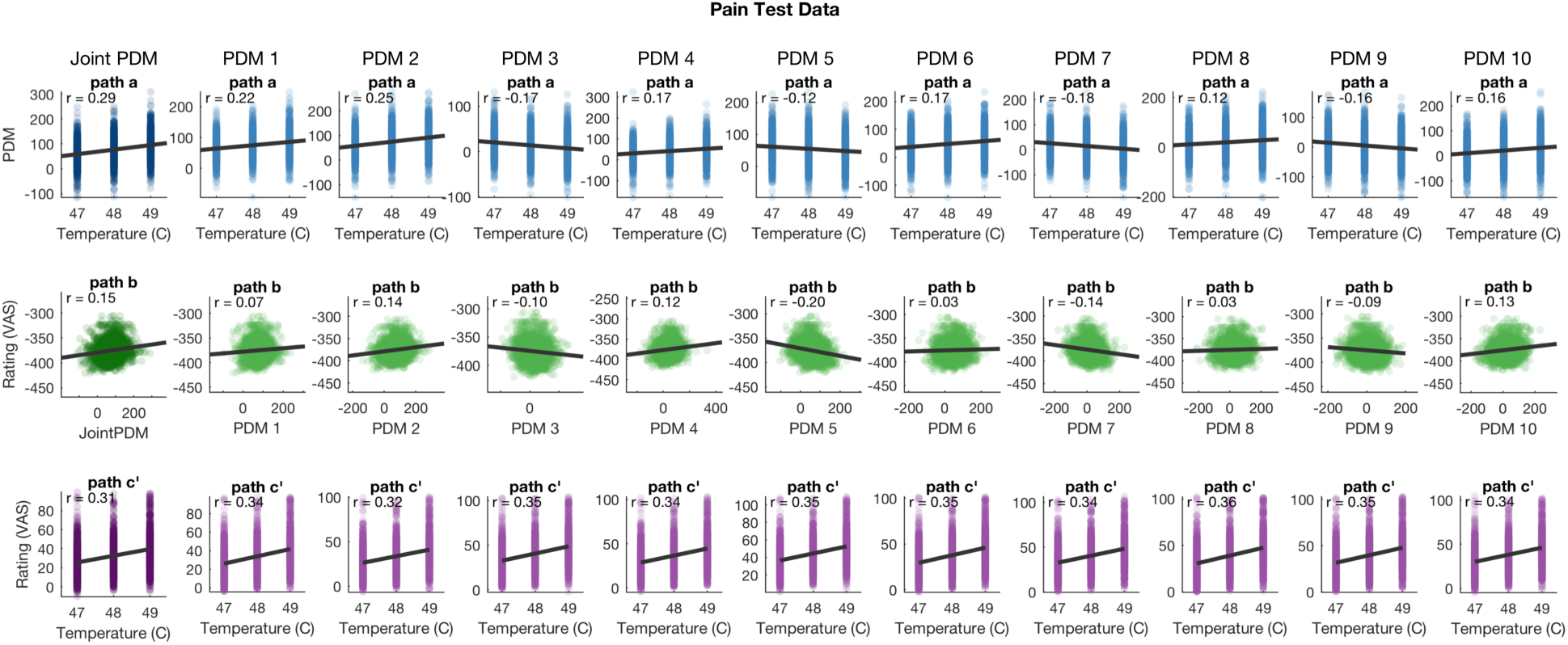
Bivariate relationships between temperatures, mediators (PDM expression), and pain ratings for the pain test data (Study 8, N = 75). Data are adjusted according to the mediation equations, i.e. ratings in path b plots are adjusted for temperatures and PDMs, ratings in path c’ plots are adjusted for PDMs, and PDMs in path b plots are adjusted for temperatures. PDMs are estimated on the pain training data (studies 1-7).

**Figure 4-supplement 2.**
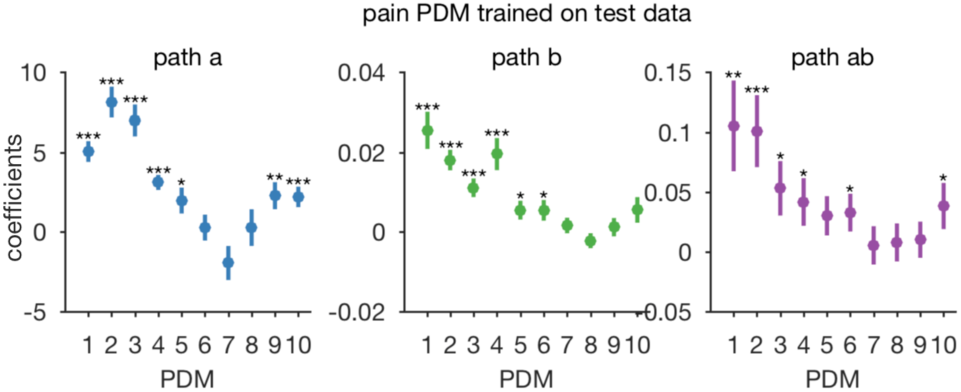
Generalization of pain PDMs from small sample to large sample. Here, test and training data sets were switched. 10 pain PDMs were estimated on the original test data set (study 8, N=75) and used as mediators in the original training data (studies 1-7, N=209). PDM 1-4, 6, and 10 are significant mediators for the larger set when trained on the smaller set.

**Figure 4-supplement 3.**
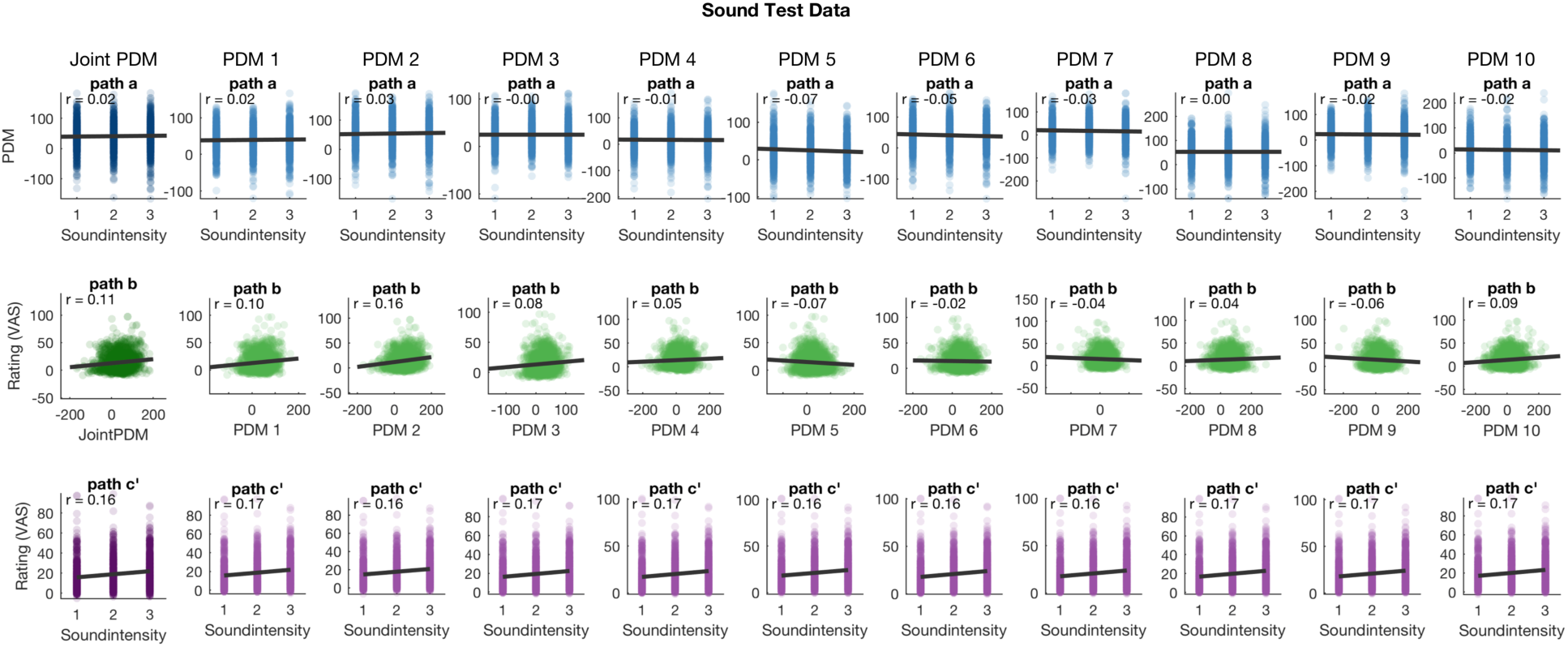
Bivariate relationships between sound intensity levels, mediators (PDM expression), and intensity ratings for the training data (studies 1-7). Data are adjusted according to the mediation equations, i.e. ratings in path b plots are adjusted for stimulus levels and PDM, ratings in path c’ plots are adjusted for PDMs, and PDMs in path b plots are adjusted for stimulus levels. PDMs are estimated on the pain training data (studies 1-7).

